# FORMAL: A model to identify organisms present in metagenomes using Monte Carlo Simulation

**DOI:** 10.1101/010801

**Authors:** Genivaldo Gueiros Z. Silva, Bas E. Dutilh, Robert A. Edwards

## Abstract

One of the major goals in metagenomics is to identify organisms present in the microbial community from a huge set of unknown DNA sequences. This profiling has valuable applications in multiple important areas of medical research such as disease diagnostics. Nevertheless, it is not a simple task, and many approaches that have been developed are slow and depend on the read length of the DNA sequences. Here we introduce an innovative and agile approach which k-mer and Monte Carlo simulation to profile and report abundant organisms present in metagenomic samples and their relative abundance without sequence length dependencies. The program was tested with a simulated metagenomes, and the results show that our approach predicts the organisms in microbial communities and their relative abundance.

## Introduction

Microbes are more abundant that any other organism (Whitman, Coleman & Wiebe, 1998), and it is important to understand what those organisms are doing and who they are. In many environments more than 99% of the microbes cannot be cultured (Sharon & Banfield, 2013).

Metagenomics is a powerful tool that makes it possible to study genomes and understand the diversity present in uncultured microorganisms just by using DNA sequences. Metagenomics uses high throughput sequencing – fast and cheap sequencing provided by the next generation of sequencing technologies (Wooley, Godzik & Friedberg, 2010).

The understanding of microbial communities is important in many areas of biology. For example, metagenomes can provide the difference the microbial community in marine animals (Trindade-Silva et al., 2012; Oliveira et al., 2012; Garcia et al., 2013).

It is not an easy task to identify the diversity of organisms present in metagenomes, and there are many approaches that have been developed to identify the organisms present in metagenomes, such as BLAST (Altschul et al., 1990), MEGAN (Huson et al., 2007), MG-RAST (Meyer et al., 2008), and Phymm (Brady & Salzberg, 2009). However, most of these tools are slow and depend on the sequence length.

Sequence composition (k-mer) analyses have been widely used to cluster unknown sequences (Rosen, Reichenberger & Rosenfeld, 2011; Silva et al., 2014), while
Monte Carlo Simulation has been previously used on high dimensional biological data (Manly, 2006; Kerr et al., 2008; Liu, 2008). However, these two approaches have not previously been combined to explore the diversity of microorganisms in different communities.

This paper proposes an approach based in a stochastic model to identify the organisms present in metagenomic samples and their relative abundance using Monte Carlo simulation and k-mer frequency from known microbial genomes.

## Methods

### Calculating k-mer frequency

Given a DNA sequence *S*, k-mer counting is a problem, in 4*^k^* - dimensional space of determining the occurrence of substrings of length k. A metagenome is a finite representation of the members of a community. The community can be considered to be a sample from all microbes. Since we are only concerned with identifying microbes that we know about, we limit the definition of community to be a sample of all *known* microbes (our reference set). Thus, k-mers could be used to predict the organisms that are present in a metagenome: the goal is to match up the organisms in the metagenome from reference dataset. There are several agile tools to count k-mers, and here we use Jellyfish (Marçais & Kingsford, 2011).

### Reference dataset

A group of reference sequences are required to model and identify the organisms that are present in a metagenome. 2,518 complete genomes were downloaded from NCBI (ftp://ftp.ncbi.nih.gov/genomes/Bacteria) on 16 November 2013, and we manually selected 1 genome to represent each genus; a total of 655 genera. K-mer frequencies (k=6-8, default: k=7) were calculated and normalized by the sum of frequencies.

### Monte Carlo Simulation

Monte Carlo methods are used as an alternative to understand the behavior in high dimensional random samples (Mooney, 1997). This method is implemented in a computer program by defining a pseudo-population under conditions from the real world. For example, we consider metagenomes as the real problem world where *M* is a random metagenome with n organisms defined as the convex combination.

We present in the sub-section “stochastic modeling” the pipeline to profile a metagenome using Monte Carlo methods.

### Distributions for biological populations

As used in PHACCS (Angly et al., 2005), our tool assumes three biological rank-abundance distributions: power (zipf) law, exponential, and log-normal distributions. The first two forms are empirical distributions used to model an asymptotic drop-off in the abundance (Ulrich, 2001) and the last form appears to be the most commonly present among species rank distribution (Pielou, 1975).

### Simulated data

The program was tested using simulated data based in complete bacterial genomes (Table 1). In order to evaluate our approach performance, a medium complexity dataset, 507,808 single 100 nt reads using the supplied error model for pyrosequencing, was sampled from ten species using Grinder (Angly et al., 2012). The ten selected species are not present into the reference dataset.

**Table 1.** – simulated metagenome.

| NCBI accession number | Strain name | Relative abundance (%) |
| --- | --- | --- |
| NC_021003.1 | *Streptococcus pneumoniae* SPN032672 | 22.85 |
| NC_016842.1 | *Treponema pallidum* SamoaD | 16.30 |
| NC_015758.1 | *Mycobacterium africanum* GM041182 | 15.09 |
| NC_015566.1 | *Serratia* AS12 | 10.55 |
| NC_009455.1 | *Dehalococcoides* BAV1 | 8.11 |
| NC_018887.1 | *Borrelia afzelii* HLJ01 | 7.33 |
| NC_020411.1 | *Hydrogenobaculum* HO | 7.11 |
| NC_008600.1 | *Bacillus thuringiensis* Al Hakam | 5.86 |
| NC_008369.1 | *Francisella tularensis* holarctica OSU18 | 4.92 |
| NC_015866.1 | *Rickettsia heilongjiangensis* 054 | 1.87 |

### Stochastic modeling

To model the data the program follows the pipeline depicted below

1. **K-mer frequency for the input data:** a script written in Python language calculates the k-mer frequency for the user input data.
2. **Select G genomes:** the algorithm selects G genomes randomly among the 655 reference genomes.
3. **Approximate distribution:** calculate the approximate distribution of the genomes in metagenomes using k-mer frequencies applies a biological distribution on the k-mer frequency of each genome and create a single vector with the sum of G genomes k-mer frequencies.
4. **Frequencies distance:** calculate the distance between the G genomes selected and the metagenome using the Euclidean distance.
5. **Repeat:** loop N times on steps 2, 3 and 4.

## Results and discussions

In order to test the tool accuracy, species from the reference dataset were not included in the simulated test set, and the genera were predicted for all the species.

For the propose of evaluating the program, it was initially run 10 times, each time with 250 x 10^4^ iterations, from 6 to 8 mers (see Figure 2).

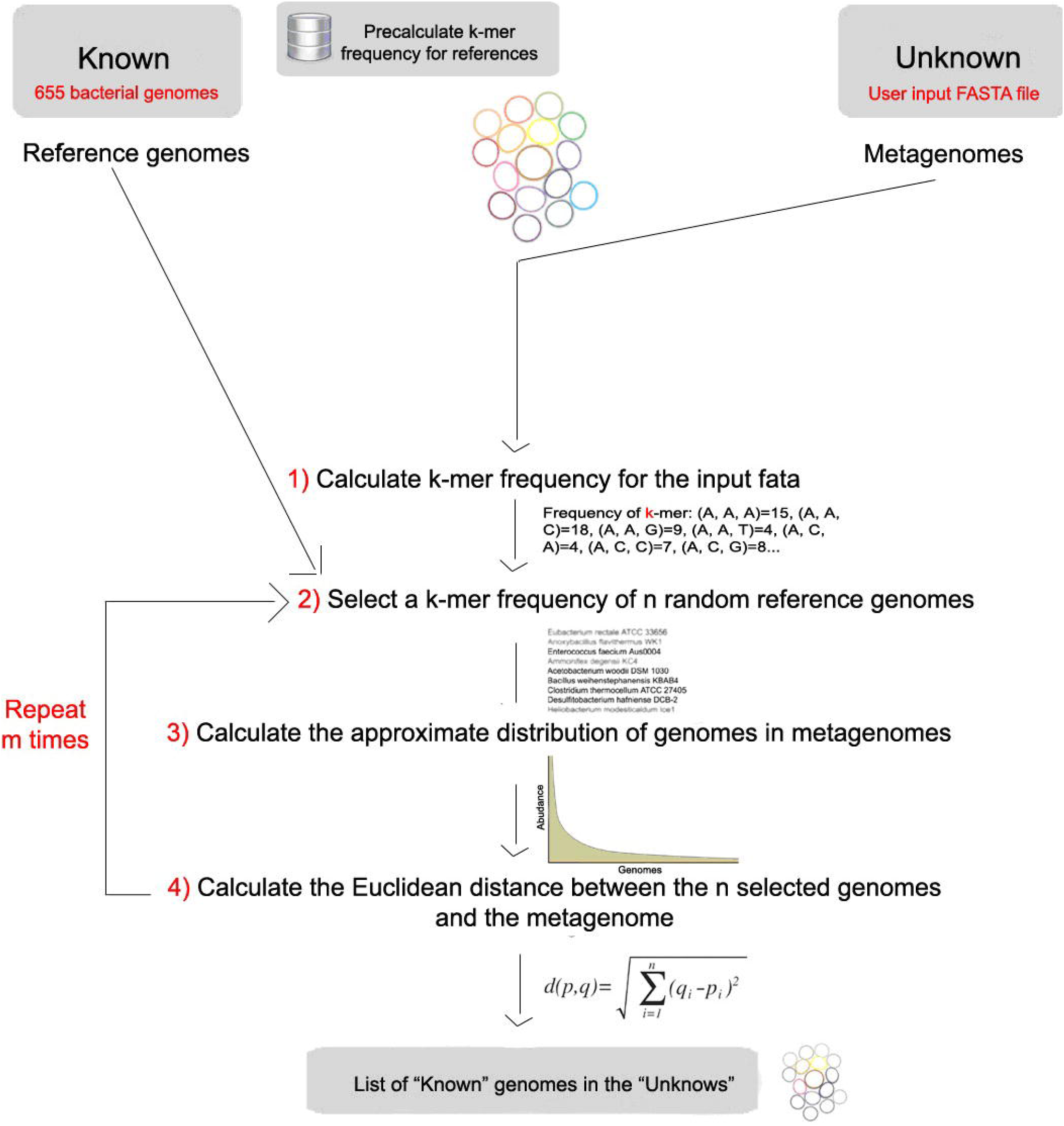

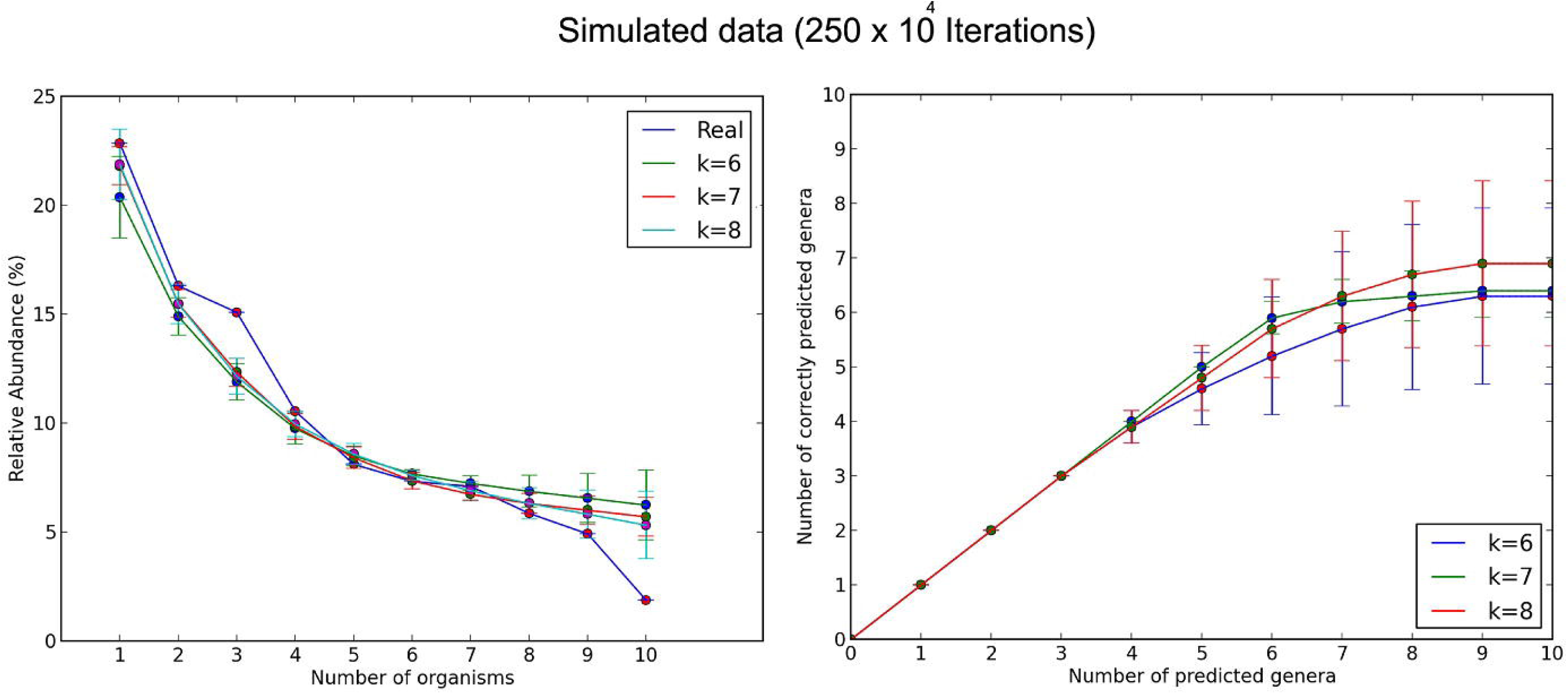

Next, I ran the program with the same parameters, but now with 250 × 10^5^ iterations (see Figure 3).

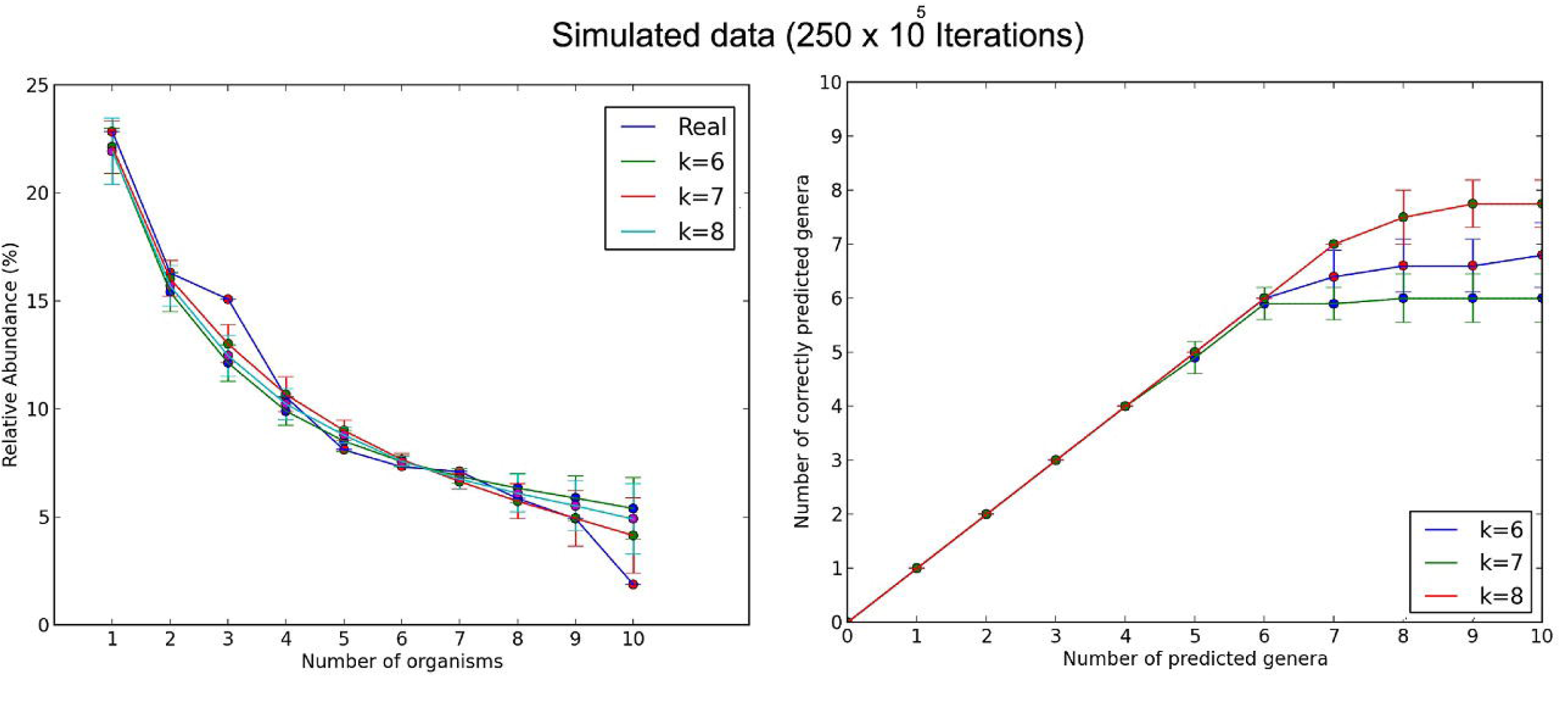

Modeling with 250×10^4^ and 250×10^5^ iterations, the stochastic approach predicted about seven or eight of the ten species among the all species in the metagenome. We clearly see in Figures 2 and 3 that when we increased the length of k, the estimation of real abundance and number of correctly predicted organisms increased. I also conclude that the error in the prediction decreases as the number of iterations increases; however, the program running time increases as the number of iterations increases (see Figure 4).

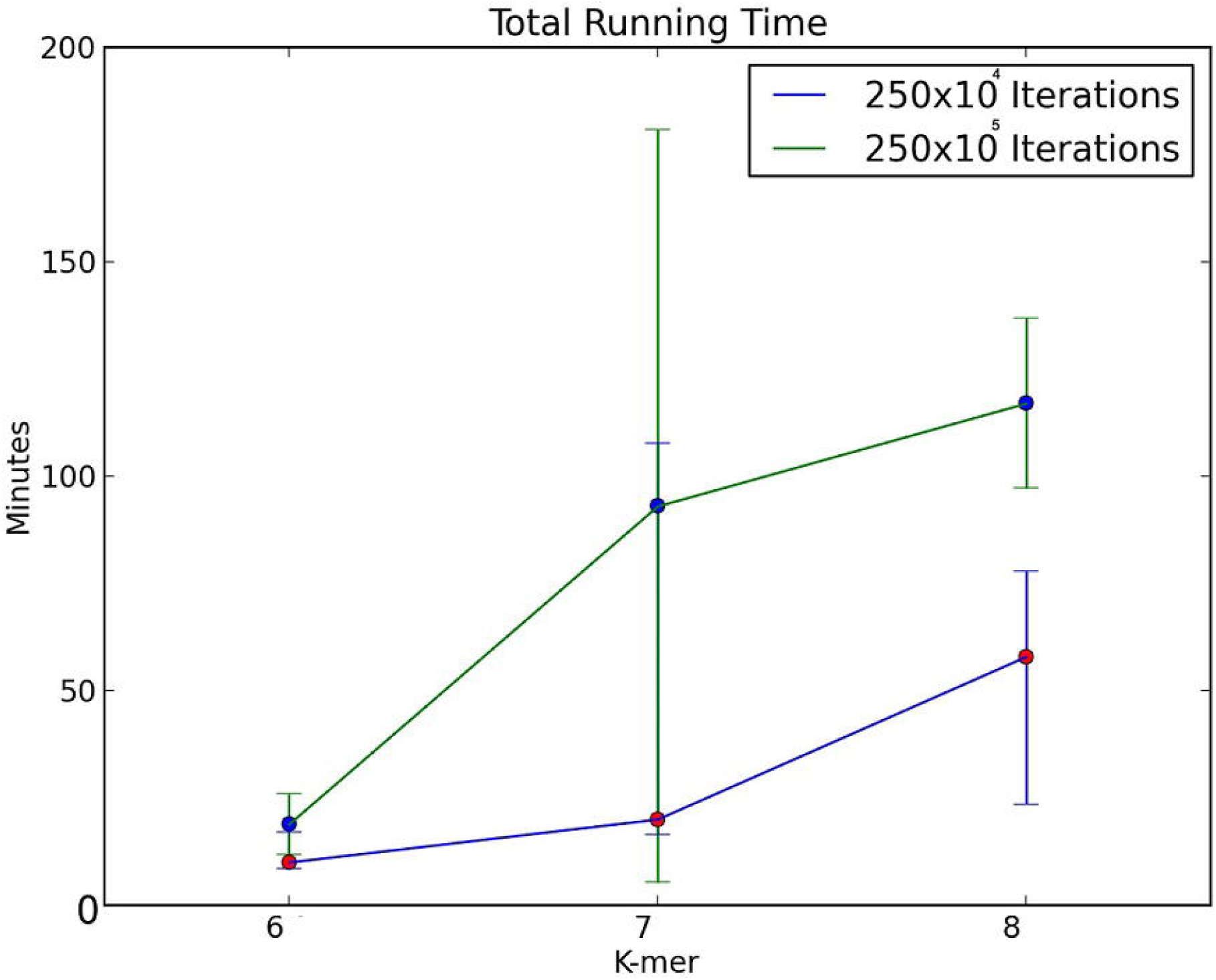

About two species were not predicted with neither 250x10^4^ nor 250x10^5^ iterations, this is probably because either the low abundance of each species in the metagenomic sequences or the presence of other species with similar k-mer frequencies in the reference database.

## Limitations

As with other methods created to profile metagenome sequences, the approach presented here depends on a curated database of microbial reference genomes in order to predict a specific genus. If a reference genome is absent, the tool will predict the closest reference available.

## Conclusions

Here a stochastic model was presented to address the problem of identifying organisms present in metagenomes that does not rely on sequence length. Thus, it can be used on raw sequencing reads and does not require an initial metagenome assembly, which may be difficult for cases with high microdiversity, or computationally costly for very large metagenomic datasets.

## Acknowledgements

I thank Dr Samuel Shen for the Mathematical modeling classes.

*Funding:* This research was funded by the Dimensions: Shedding Light on Viral Dark Matter project supported by the National Science Foundation (NSF) Division of Environmental Biology (DEB-1046413) and by II-EN: Computational Enhancement of Analytical Metagenomics Systems project supported by the NSF Division of Computer and Network Systems (CNS-1305112).

*Conflict of Interest*: none declared.

### Availability and requirements

Project home page: https://edwards.sdsu.edu/formal

Operating system: the program was developed for Linux but should also run on Windows or Mac command line interpreters (Cygwin, Terminal).

Programming language: Python.

